# GLACIAL SPECIALIST SPECIES ARE MORE VULNERABLE TO CLIMATE CHANGE THAN GENERALIST SPECIES (DRAWING PARELLELS BETWEEN EMPIRICAL AND ECOLOGICAL DISTINCTIONS)

**DOI:** 10.64898/2026.09.04.749327

**Authors:** Bart Steen, Alejandra Morán-Ordóñez, Antoine Guisan, Gianalberto Losapio

**Author notes:** Corresponding authors: Bart Steen:, Gianalberto Losapio. Shared last co-authorship Institutions (Antoine Guisan, Gianalberto Losapio). Author contact info: Alejandra Morán-Ordóñez:; Antoine Guisan.

## Abstract

In Switzerland, glaciers are projected to shrink to minimal size by the end of the century. Thus, climate change puts pressure on glacial specialist plant species, as generalist plant species colonise their habitat. As bare areas are revealed by glacier retreat, glacial specialists can likewise shift their distributions to higher altitude, however, there will be no habitat for them to colonise. Thus, Swiss mountain floral biodiversity and ecosystem services will likely be locally extirpated. In this article, we use Species Distribution Models of 369 plants occurring in Swiss proglacial areas fitted using bioclimatic, lithological and topographic variables and compare the covariates that most strongly predict the distributions of glacial specialist and generalist species. In addition, we use two methods of defining glacial specialist species and separating them from specialists: empirical (degree of prevalence of species in proglacial areas) and ecological (using Landolt values). We find that both approaches yield similar results: the distributions of both generalist and glacial specialist species are most strongly predicted by bioclimatic variables, however, they are significantly stronger for glacial specialists. On the other hand, distributions of generalist species are more strongly predicted by topographical and lithological variables.

## INTRODUCTION

The Swiss glaciers are predicted to shrink to minute size by the end of the 21st century (Zekollari et al. 2019). Their rapid recession is exposing bare substrate that is being colonized by glacial specialist species, and older colonized areas are subject to ecological succession (Charles et al. 2025, Khelidj et al. 2026). This will lead to a temporary local increase in mountain biodiversity, as the intermediate stage of succession is the richest in biodiversity (Cannone et al. 2022, Tu et al. 2025). However, the climax stage is poorest in biodiversity, consisting mostly of generalist species (Charles et al. 2025). In addition, there will eventually be no more area for glacial specialists to colonize. Thus, succession in proglacial areas jeopardizes the survival of unique mountain flora, as well as the ecosystem services and functions they provide (Losapio et al. 2025), and eventually, even the dynamic intermediate stage will make way for the climax stage (Ficetola et al. 2021, Cannone et al. 2022, Mainetti et al. 2022, Tu et al. 2025). As such, there will be clear “winners and losers”, with generalists most likely to thrive and glacial specialists most at risk (Rumpf et al. 2018).

In Switzerland, the shifting glacier forelands will be characterized by highly localized changes in environmental conditions, such as substrate pH (as soils form and change) (Charles et al. 2025), microclimatic variations, and microtopographic heterogeneity (Scherrer & Körner 2011), but also by broader environmental changes, such as increasing precipitation seasonality, rising mean annual temperature (MeteoSwiss & ETH Zurich 2025) and topographic changes, which include increased steepness of slopes (Savi et al. 2021). If glacial specialists are more sensitive to these combined pressures, which is probably the case (Rumpf et al. 2018), they should warrant higher conservation priority. Climax species, on the other hand, might benefit from these changes if they are less sensitive to them. Scientific knowledge on this issue is fragmented, and we lack broad-scale knowledge of which species are more vulnerable to each risk factor, as well as of which risk factors are the strongest determinants. A possible reason for this is the lack of a unified standardized guideline and framework.

Species Distribution Models (SDMs) emerged in the last three decades as a powerful tool to assess species-environment relationships and hence to infer the sensitivity of species to environmental conditions (Guisan et al. 2017). In addition, SDMs can be used to make standardized applicable decisions in conservation prioritization by guiding the allocation of limited available resources (Steen et al. 2019, Steen et al. 2024, Guisan et al. 2025). Yet, considering the ongoing and future environmental changes in proglacial areas, we lack also a general approach to distinguish between glacial specialists and generalists. In this study, we address this paucity by presenting a framework that combines an empirical and an ecological distinction between glacial specialist species and generalist species. We first used SDMs of plant species present in Swiss proglacial areas to compare the explanatory power of different covariates associated with the environmental changes in proglacial areas between the two species groups, distinguished by different measures. The empirical distinction was made using the geographic locations of the known occurrences of the modelled species inside and outside proglacial areas; the latter was determined using Landolt values of the modelled species, which is independent of SDM output. This may provide a generalized and transferable method for assessing the modelled species’ specialization and, hence, their vulnerability.

We researched this by addressing two research questions: (1) Which species are most sensitive to which environmental changes, and how sensitive are they? This can be quantitatively assessed using the SDMs. (2) Which environmental drivers represent the most critical risk factors shaping species responses in Swiss proglacial areas?

We hypothesize that glacial specialists are more sensitive to changes in environmental conditions than generalists, responding both more frequently and more strongly to the risk factors associated with glacier retreat.

## METHODS

### Overview

Occurrence records of 721 vascular plants species from the Infospecies organization inside Swiss proglacial areas were used. These areas were defined as the regions in Switzerland deglaciated since 1850, as determined by GLAMOS (Glaciers monitoring Switzerland, n.d.). Of these, species that either had ten occurrence records in proglacial areas, or a relative prevalence of 1% in proglacial areas (meaning 1% or more of the InfoSpecies occurrence data found themselves in proglacial areas), or both, were selected, leaving 390 species.

Subsequently, occurrence records form GBIF were collected for all 390 species (GBIF Occurrence Download 2023) and used in a spatially-nested Species Distribution Modelling (N-SDM) approach (Guisan et al. 2025). Models were fitted at European scale using bioclimatic variables, which in turn were used as an input variable for a model fitted at Swiss scale, which was fitted additionally using topographic, lithological and local bioclimatic variables. For a list of the variables used to fit the models, see Table 1. The optimal models were projected at Swiss level.

**Table 1:**
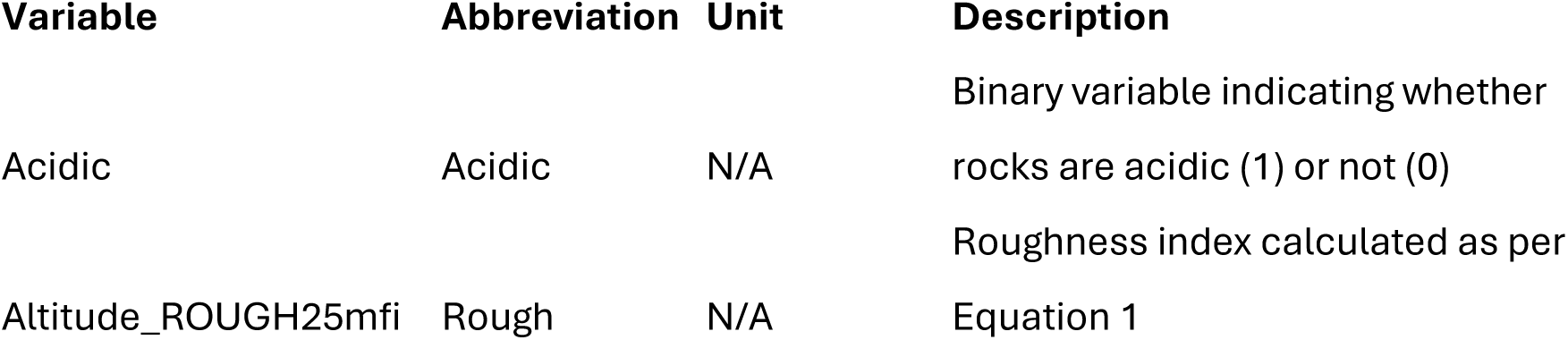
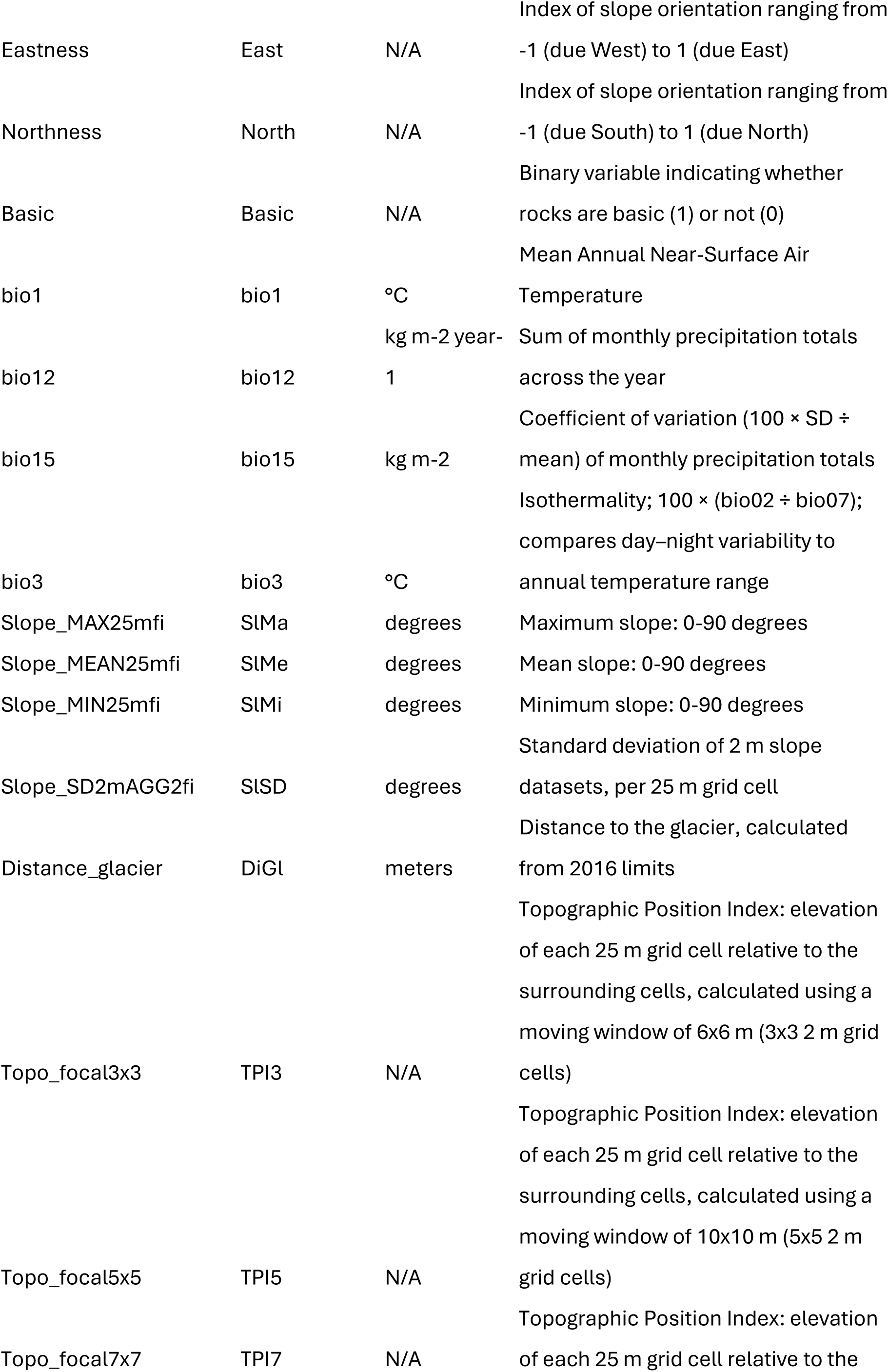
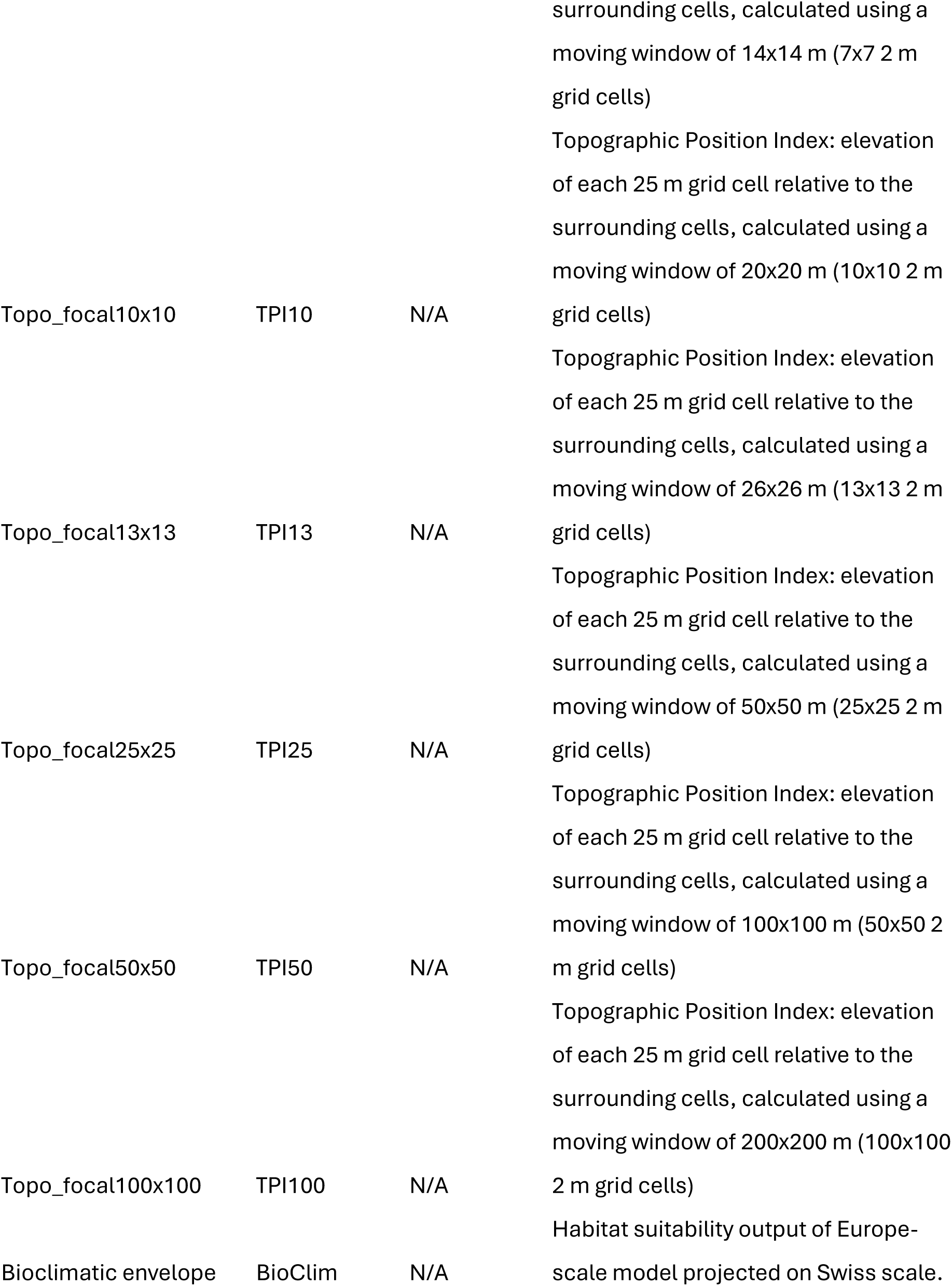
Names, abbreviations, units and descriptions of all variables used in the Swiss-scale SDMs. The resolution for all variables was 25×25 m and the coordinate reference system was EPSG:205c (CH 10S3+ /LVS5).

21 Swiss-scale models returned invalid performance metrics, leaving 369 modelled species. For the final list of modelled species, see Table 2. Our study species contained 198 hemicryptophytes, 69 chamaephytes and 8 phanerophytes (including nanophanerophytes); see Supplementary materials, Table S3.

For an overview of which species were included and why they were or were not kept for modelling, see supplementary materials, table S1.

Optimal models for each species were selected according to the maximum True Skill Statistic (maxTSS) value. Subsequently, the modelled species were divided into glacial specialists and generalists. This was done in two ways: as per the prevalence level of occurrence records in proglacial areas and according to Landolt ecological values (Landolt et al. 2010), forming an empirical and an ecological approach, respectively.

Of the models run for the two species groups, the average variable importance of all variables of the models at Swiss scale (which included the model output of the Europe-scale SDM) were extracted and the average results among the two species groups were compared.

For an overview of the project workflow, see Figure 1.

**Figure 1:**
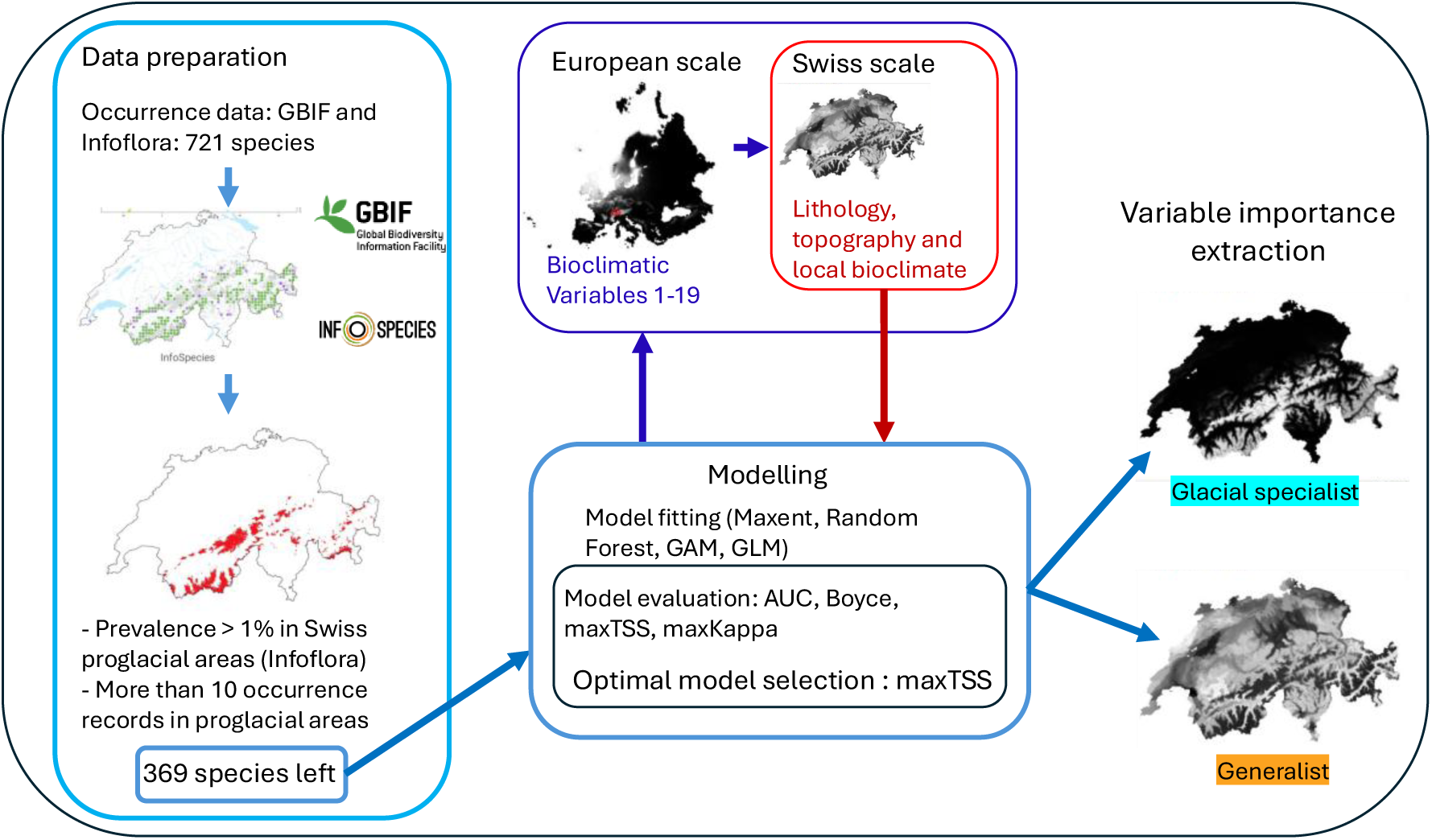
Design of the study. Notice the nested approach: the output of the European models is used as an input variable for the Swiss-scale models, which are in turn fitted, the best model is chosen based on the highest maxTSS value, and the variable importance is finally extracted. This was done for all 3cS species.

### Occurrence data

From the data from Infospecies of species with at least 10 occurrence records inside proglacial areas, the data with less than 25m resolution were removed. For the Europe-scale bioclimatic models, occurrence data was downloaded from GBIF from between the years 1980 and 2023 (GBIF occurrence download 2023) for all the species identified by the Infospecies data. The species names were checked for synonymous names of the same taxon and if these were found, occurrence data bearing those names were downloaded, as well, in order to avoid mismatching and losing data. Duplicated records, data with an unknown taxon rank and data with a coordinate uncertainty of more than 1000m were removed.

### Species Distribution Modelling workflow

A hierarchical approach (Guisan et al. 2025) was used to fit SDMs for 369 species. The occurrence data from GBIF and of Infospecies were used to fit the Europe-scale models (climatic niche; bioclimatic envelope), during which only one occurrence point per grid cell (meaning per 30 arc seconds) was used. This model was subsequently projected at Swiss level, at 25m resolution, by projecting the bioclimatic variables on the matching variables of the SWECO database (Külling et al. 2024). Thus, the bioclimatic envelope was used as an input variable to fit the Swiss-scale SDM, which trained using the Infospecies data (not the GBIF occurrences). Again, only one occurrence point per grid cell, in this case per 25m, was used to train the Swiss-scale models. All models were fitted using 10000 background points sampled randomly in geographic space.

At both scales, the covsel R package (Adde et al. 2023) was used to automatically select variables from the respective candidate lists, meaning that the covariates for each model were chosen from a parsimonious set that explained the highest possible amount of variation in the species’ distributions.

Models were fitted using Biomod2 (Thuiller et al. 2009), using four different algorithms: MaxEnt (the maxnet version (Phillips et al. 2017)), Generalized Linear Models (GAM), Generalized Additive Models (GAM) and Generalized Boosted Regression Models (GBM). Their performance was quantified using the Kappa, True Skills Statistic (TSS) and Area Under Curve (AUC) metrics of test datasets obtained from random partitioning of the occurrence data (80% training and 20% test). The same number of models was run for each species. Finally, the BIOMOD_EnsembleModelling() function was used to combine the models with the highest TSS score into one. For two Swiss-scale model projections of example ensemble models, see Figure 2.

**Figure 2:**
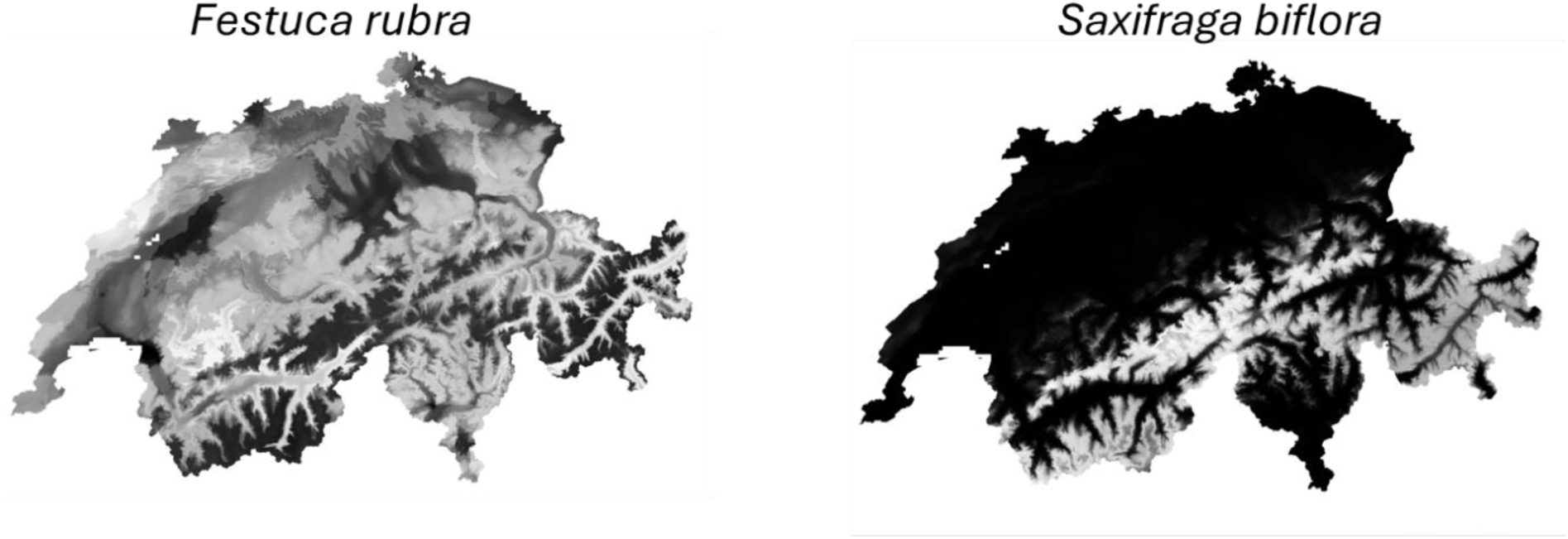
examples of projected maps of final models fitted at Swiss scale for a generalist species (Festuca rubra) and a glacial specialist species (Saxifraga biflora).

### Environmental variable creation and selection

For the the Europe-scale models, the 19 CHELSA V.2.1 bioclimatic variables at 30 arc second resolution (Karger et al. 2017) were used to fit the model at Europe scale, making it the species’ bioclimatic envelope.

For the Swiss-scale model, environmental variable maps were constructed as raster layers at a 25 m resolution, covering the extent of Switzerland. Topographic slope angle (2 m resolution, Swiss Federal Office of Topography) was aggregated using ArcGIS Pro (ESRI 2024) to derive four slope angle metrics: mean, minimum, and maximum. The original 2 m dataset was first aggregated to 26 m resolution and subsequently resampled to 25 m using nearest-neighbor interpolation to match the spatial resolution of other environmental variables. Standard deviation of slope angle was also calculated from the 2 m DEM. Similarly, mean elevation was derived from the same 2 m resolution Digital Elevation Model (DEM). The Topographic Position Index (TPI), quantifying topographic concavity-convexity, was calculated using moving windows (3×3, 5×5, 7×7, 13×13, 25×25, 50×50 and 100×100 cells) applied to the DEM. In addition, two measures of terrain roughness were generated: a standard deviation of slope (13 x 13 cells, resampled to 25 m) and a Topographic Roughness Index calculated using ArcGIS, applied to each 25 m grid cell (formula 1).

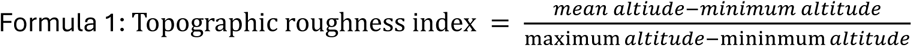

Aspect orientation was quantified from a 2 m resolution aspect dataset as northness (degrees North-facing) and eastness (degrees East-facing), with values ranging from 0 to 360 degrees. Northness and eastness were derived using sine and cosine transformations, respectively (Guisan et al., 1999), resulting in values from 1 (North/East) to −1 (South/West). Four bioclimatic variables from the SWECO25 dataset (Külling et al., 2024) were included: annual mean temperature (bio1), isothermality (bio3), annual precipitation (bio12), and precipitation seasonality (bio15). These variables, at 25 m resolution and validated against local weather station data, were selected for their low correlation within the Swiss region and interpretability. Finally, datasets representing alkaline and acidic bedrock types from the Swiss Federal Office of Statistics (OFS, Maps of Switzerland, n.d.) were incorporated. The shapefiles were used to extract those areas on the 2m DEM layer that contained acidic substrate types, the cells within and outside the areas were given a value of 100 and 0, respectively, and the resulting binary map was aggregated from 2 m to 26 m resolution using the sum criterium, after which it was resampled to 25m. The same was done for the basic map. Due to the aggregation, the maps contain values between 100 and 0 at the edges between acidic/basic and non-acidic/basic. A comprehensive list of all variables used to fit the SDMs is given in Table 1.

### Species threshold variation

Several thresholds of occurrence prevalence to draw the distinction between glacial specialists and generalists were utilized: firstly, the empirical prevalence threshold, which is the threshold of the minimum percentage of occurrence records obtained from InfoSpecies that find themselves in proglacial areas. Secondly, the ecological threshold, derived from Landolt values, which were the species belonging to the species belonging the indices Temperature, class 1 (= plants belonging to nival and glacial habitats), Light index class 5 (= plants intolerant of shade), and Moisture variability index class 1 (minimal moisture variation).

Then, for each distinction criterium, the average variable importance of all models for glacial specialists and for generalists (so not the ensemble models) were compared with an unpaired t-test (Student’s or Welch’s, depending on whether equal variances could or could not be assumed, respectively). This was done using the “t.test” and “var.test” functions in R (R core team, 2021).

The variables were pooled into three general groups: bioclimate (including the Europe-scale output), lithology and topography. We also did tests that separated the Europe-scale model output from the other bioclimatic variables.

## RESULTS

See figure 2 for the ensemble model projections of two species that were always respectively included in one of the two groups – *Festuca rubra*, a generalist, and *Saxifraga biflora*, a glacial specialist.

### Species Distribution Model performance metrics

For the bioclimatic envelope, meaning the Europe-scale models of the 369 species, the average Kappa was 0.591 (±0.102), average TSS was 0.686 (±0.069) and average AUC was 0.915 (±0.032). Maximum and minimum Kappa were 0.731 and 0.328, respectively, maximum and minimum TSS were 0.796 and 0.504, respectively, and maximum and minimum AUC were 0.95 and 0.829, respectively.

For the Swiss-scale models, average Kappa was 0.533 (±0.056), average TSS was 0.722 (±0.053) and average AUC was 0.931 (±0.018). Maximum and minimum Kappa were 0.684 and 0.421, respectively, maximum and minimum TSS were 0.835 and 0.606, respectively, and maximum and minimum AUC were 0.971 and 0.891, respectively. Overall, this indicates a good capacity of the models to discern between suitable and unsuitable habitat for the species.

### Empirical approach variable importance

See Figure 3 for a summary of the main results of empirical separation between generalists and glacial specialists. The average variable importance of the bioclimatic variables differed significantly between generalist species and glacial specialists, with the latter being more powerfully predicted as such. At 5% prevalence threshold, the glacial specialists’ average importance of bioclimatic variables was 0.262 (n = 102), versus 0.211 (n = 267) of the generalist species (t-test: p = 3.483e-10). The pattern did not significantly change when the prevalence threshold was varied. Even when placing the limit at 20%, which left only 15 glacial specialist species, the difference was still significant (t-test: p = 0.016).

**Figure 3:**
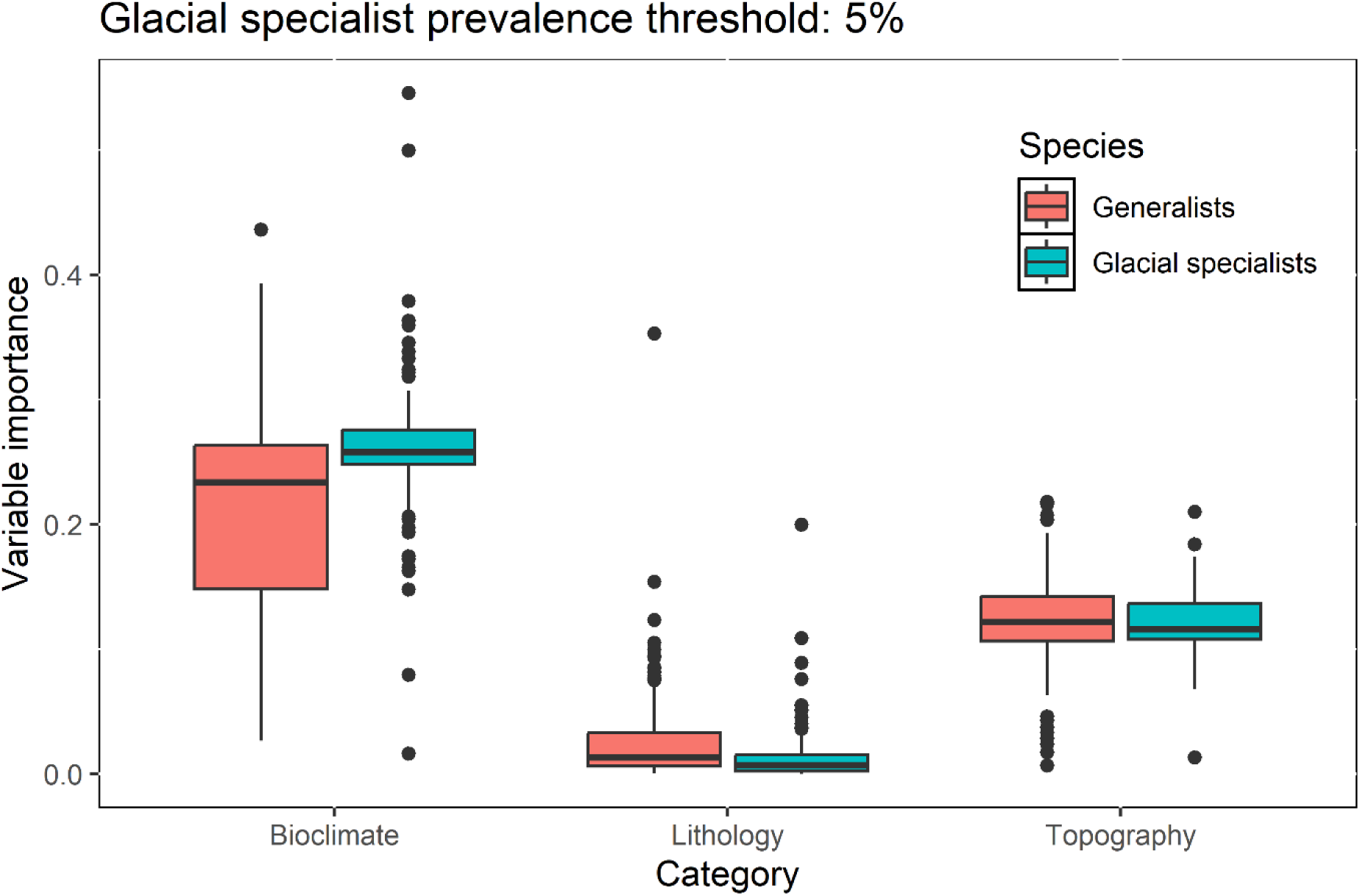
Average variable importance of the bioclimatic, lithology and topographic variables in the SDMs for glacial specialists versus generalist species. The distinction between glacial specialists and generalists was drawn by setting the prevalence threshold of the glacial specialists in proglacial areas to 5%.

The lion’s share of the variable importance was carried by the model output at Europe level, i.e., the bioclimatic envelope. At 5% prevalence threshold, the average variable importance was 0.85 for glacial specialist species and 0.61 for generalist species (t-test: p = 6.226e-16).

Conversely, local bioclimate yielded significantly higher average variable importance for generalist species (0.072) than for glacial specialists (0.057) (t-test: p = 0.0479). The 5% prevalence however was the only empirical threshold for which a significant effect could be detected. For a full overview of the variable importances per prevalence threshold, see supplementary materials, table S1.

Lithological variables had higher average variable importance for generalist species (0.024) than for glacial specialists (0.015) (t-test: p = 0.004). Topographical variables never yielded significant results at any prevalence threshold, even though the average variable importance is relatively high (0.121 for glacial specialists and 0.125 for generalists).

### Ecological approach variable importance

See Figure 4 for a summary of the main results of ecological separation between generalists and glacial specialists. They were obtained from the T_index = 1, meaning the glacial specialist category contained only species belonging to nival and glacial habitats according to Landolt (2010). This was the only ecological separation where the average variable importance of all bioclimatic variables put together (meaning they included both the bioclimatic envelope and the local bioclimatic variables) for glacial specialists were significantly higher (0.268, n = 87) than for generalists (0.212, n = 282) (t-test: p < 2.2e-16). The average variable importance of the bioclimatic envelope only was always significantly higher for glacial specialists than for generalists, no matter which ecological distinction as used (T-index = 1, L-index = 5 or W-index = 1) and local bioclimate variable importance was significantly higher for generalists than for glacial specialists only for the T-index variation. See supplementary materials, table S1 for details.

**Figure 4:**
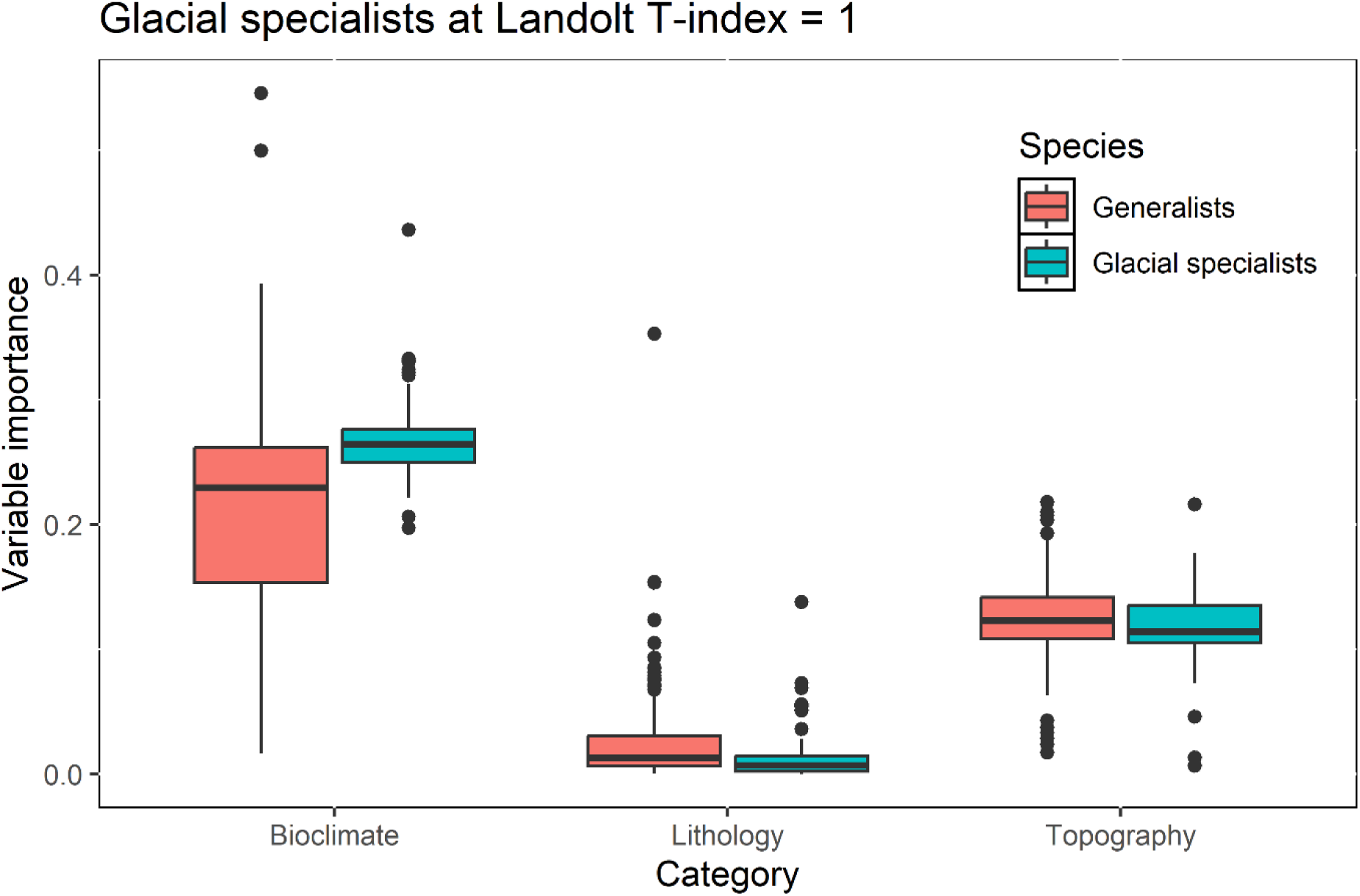
Average variable importance of the bioclimatic, lithology and topographic variables in the SDMs for glacial specialists versus generalist species. The distinction between glacial specialists and generalists was drawn by selecting those modelled species that had a Landolt T-index of 1, meaning those species that (in Switzerland) belong to glacial and/or nival habitats.

Topography was a significantly more powerful predictor for generalists than for glacial specialists (0.125 vs 0.118, t-test: p = 0.04248). The same holds true for lithology (0.024 vs 0.013, t-test: p = 0.0003).

### Variable selection frequency

The average frequency of selection was calculated as follows: first, we determined how often a variable belonging to each category was selected. This means that the number of models in which any variable of the chosen categories (local bioclimate, lithology and topography) for each species was recorded. Then, these numbers were summed for each variable category for both glacial specialists and generalists, and these final respective values were divided by the number of species in each category. The bioclimatic envelope was always forced to be in the models, and thus, 100% of all models of both glacial specialists and generalists contained that variable. For local bioclimate, the average frequency was 11.6 for glacial specialists versus 12.0 for generalists (t-test: p = 0.03012) when separating by 5% prevalence. For lithology, it was 7.8 versus 7.9 (t-test: p = 0.3358) and for topography, 20.8 vs 20.9 (t-test : p = 0.824).

For the T-index variation, the average values of glacial specialists vs. generalists were 11.7 vs 12.0 (t-test: p = 0.1371) for local bioclimate, 7.9 vs 7.9 for lithology (t-test: p = 0.770) and 20.2 vs 21.0 for topography (t-test: p = 0.036).

See supplementary materials, figure S5 for the variable selection results at 5% prevalence and T-index = 1.

## Discussion

As the Alps will see significant changes in climate, and species succession will affect areas where pioneers have been established (Ficetola et al., 2024; Tu et al., 2024, Charles et al. 2025) altering the vegetation composition and likely favouring generalist species whilst jeopardizing glacial specialists and their associated ecosystem functions (Losapio 2025). This study aimed to confirm if the environmental changes taking place in the Alps will indeed disproportionately affect glacial specialists over generalist species (research question 1), which would be expressed both by strength and frequency of selection of the corresponding factors in the SDMs. We aimed also to determine which abiotic factors pose the greatest risk of species extirpation (research question 2).

The results indicate that for both our empirical and ecological method of distinction, glacial specialists are more sensitive than generalists to environmental changes that will take place in the Swiss Alps, most notably bioclimate. Mean annual temperature, isothermality and precipitation seasonality will all increase (MeteoSwiss & ETH Zurich 2025). These changes are covered directly in our local variables (bio1, bio3 and bio15, respectively) and indirectly in the bioclimatic envelope. All these influence glacial species significantly more than generalists. This confirms our hypothesis and is likewise confirmed by other studies, which indicate likewise that generalists are colonizing Alpine summit floras (e.g., Liberati et al. 2019). The hypothesis is also confirmed by the frequency of selection, however only for the empirical approach, though the difference is marginal. A possible reason for this is that, since the covsel R package uses statistical algorithms to select a parsimonious set of predictor variables according to the amount of variance they explain (Adde et al. 2024) and both scales of bioclimate were still often selected, as they are relatively powerful predictors (see figure 3 and 4).

As for research question 2, our results clearly show that the most significant environmental drivers in both glacial specialist and in generalist species distributions are climatic, for both our methods of determining them (ecological and empirical). This is expected, as temperature and precipitation in Alpine habitats regulate many ecological factors, like snow cover (Klein et al. 2016, Poloczanska et al. 2018, Francon et al. 2021), life-history traits (Dullinger et al. 2012, Ficetola et al. 2021), phenology (Theurillat C Schlussel 2000, Vitasse et al. 2021), etc. In addition, the bioclimatic envelope (meaning the Europe-scale model output) is a stronger predictor than the local ones.

This discrepancy in variable importance between the bioclimatic envelope and local bioclimate for glacial specialist species is potentially explained by the bioclimatic envelope by construction being forced to be included as a covariate, and having so much explanatory power that there is little variation left for the local bioclimatic variables to explain. The high explanatory power of the bioclimatic envelope has been confirmed in other studies (e.g., Steen et al. 2024). The significantly higher average variable importance of local bioclimatic variables for the generalist species as compared to glacial specialists might be explained by the bioclimatic envelope being able to draw an accurate distinction between climatic conditions in proglacial areas and more common ones, thus accurately predicting the climatic preferences of glacial specialists. In the case of generalists, the pattern is more blurred, as this species group is less affected by climate, and hence, there is more variation in species distributions left local bioclimatic variables to explain.

The significantly higher average lithological variable importance for generalists than for glacial specialists is probably, for a large part, a topographic effect; different areas in the Alps are rich in granite, silica-rich gneiss, and ophiolites (Hartman & Moorsdorf 2012, Geurden et al. 2025), which give rise to acidic soils (Böhner et al. 2008; Charles et al. 2025), whilst other areas contain many carboniferous and calcareous sediments (Strauss et al. 2023; Charles et al. 2025), which give rise to basic soils (Böhner et al. 2008). Alpine plants communities vary strongly with soil pH (Vittoz et al. 2010), and if generalists find themselves in more lithologically homogeneous areas than glacial specialists, this would make lithology a powerful predictor, as the models would be able to identify the contrast between basic and acidic substrates more easily. This may be supported by scientific findings that the Swiss alpine soils acidify as succession progresses, due to increasing C/N ratio, which is in turn caused primarily by incipient forest species (Charles et al. 2025). Our results may therefore reflect the acidification of the Alps. However, most of our study species are hemicryptophytes (198 out of 369), the second biggest group are chamaephytes (69 out of 369) and only a few are (nano)phanerophytes (8 out of 369), which may indicate that the commencing colonization by forest species is only in its early stages. Our results therefore are not conclusive in this regard. Lastly, Vittoz et al. (2010) found that in the Alpine lowlands until the treeline, basic (calcareous) soils hosted more diverse species communities than acidic (silicate) substrate, but that this effect disappears closer to the mountain summits. Whilst this is not necessarily a ubiquitous phenomenon (e.g., Erschbamer et al. 2006), it may point to a process that might explain our finding that lithology is a weaker predictor for glacial specialists, as it may blur the effect of lithology, though this process is, by our knowledge, not yet identified.

Topography being a more powerful determinant of generalists’ species distributions than for those of glacial specialists (albeit only for our ecological approach of separating glacial specialists from generalists) is possibly explained by nival habitats being very topographically heterogeneous (Carrer et al. 2019), meaning that glacial specialists would be adapted to high degrees of terrain roughness, slope angle, and concavity-convexity variation. As succession proceeds, less well-adapted generalist species are exposed to the geographical heterogeneity of high-mountain areas, which may limit their distributions. Another explanation is that the heterogeneous topography creates microhabitats that can provide refuges for plant species (e.g., Scherrer & Körner 2011, Geurden et al. 2025). This effect is however at odds with our findings, as the topographic variables were derived from a 2 m DEM which would likely capture such environmental variations, and yet, both global and local microclimate are more powerful predictors. In addition, topographical variables do not predict the distributions of either species group more powerfully for our empirical method of separating them. Our results therefore concur more with previous findings that over long time periods, microrefugia will no longer be effective to preserve Alpine species (Jiménez-Alfaro et al. 2024, Helm et al. 2024).

In conclusion, our hypothesis that glacial specialist species are more vulnerable to the environmental changes that will take place in the Alps than generalist species is fully confirmed. Both our methods of distinguishing between the two species groups yield the same results: bioclimate is a more powerful determinant of glacial specialist distributions, which ans wers research question 2, and the marked changes that will take place in the Alps will disproportionally affect these species. Topographic and lithological variables are generally more powerful predictors for generalists, but for both species groups, they explain considerably less variation in species distributions than bioclimate does. Lastly, to finish answering out second research question, global bioclimate drives species distributions more strongly than local bioclimate, though the effect is largest for glacial specialists.

## ACKNOWLEDGEMENTS

The study was financially supported by the Biodiversa+ PrioritIce project (Project number: 31BD30_209629; grant agreement no. 101052342). GL was also supported by the Swiss National Science Foundation (PZ00P3 202127) and the Next Generation EU, Italian Ministry of Research and Education (PRIN 2022 PNRR P2022N5KYJ).

