## Supplementary materials for "GLACIAL SPECIALIST SPECIES ARE MORE VULNERABLE TO CLIMATE CHANGE THAN GENERALIST SPECIES (DRAWING PARELLELS BETWEEN EMPIRICAL AND ECOLOGICAL DISTINCTIONS)"

***Table S1:*** *Average variable importance of the bioclimatic, lithology and topographic variables in the SDMs for glacial specialists versus generalist species and matching results of unpaired t-tests. Local bioclimate (Swiss scale) and global bioclimate (Europe scale) data are shown separately. The distinction between glacial specialists and generalists was drawn by varying the prevalence threshold of the glacial specialists in proglacial areas.*

A. Global bioclimate

| Prevalence | >=20% | >=10% | >=5% | >=1% |
|---|---|---|---|---|
| Number of glacial specialist species | 15 | 46 | 102 | 279 |
| Number of generalist species | 354 | 323 | 267 | 90 |
| Average variable importance glacial specialists | 0.83 | 0.824 | 0.85 | 0.81 |
| Average variable importance generalists | 0.67 | 0.654 | 0.61 | 0.43 |
| Unpaired t-test | p = 0.097 | p = 7.479e-05 | p = 6.226e-16 | p < 2.2e-16 |

B. Regional bioclimate

| Prevalence | >=20% | >=10% | >=5% | >=1% |
|---|---|---|---|---|
| Number of glacial specialist species | 15 | 46 | 102 | 279 |
| Number of generalist species | 354 | 323 | 267 | 90 |
| Average variable importance glacial specialists | 0.073 | 0.064 | 0.057 | 0.068 |
| Average variable importance generalists | 0.068 | 0.069 | 0.072 | 0.069 |
| Unpaired t-test | p = 0.760 | p = 0.635 | p = 0.0479 | p = 0. 8848 |

C. Lithology

| Prevalence | >=20% | >=10% | >=5% | >=1% |
|---|---|---|---|---|
| Number of glacial specialist species | 15 | 46 | 102 | 279 |
| Number of generalist species | 354 | 323 | 267 | 90 |
| Average variable importance glacial specialists | 0.013 | 0.011 | 0.015 | 0.023 |
| Average variable importance generalists | 0.023 | 0.023 | 0.024 | 0.017 |
| Unpaired t-test | p = 0.217 | p = 0.0001 | p = 0.004 | p = 0.07 |

D. Topography

| Prevalence | >=20% | >=10% | >=5% | >=1% |
|---|---|---|---|---|
| Number of glacial specialist species | 15 | 46 | 102 | 279 |
| Number of generalist species | 354 | 323 | 267 | 90 |
| Average variable importance glacial specialists | 0.124 | 0.119 | 0.121 | 0.122 |
| Average variable importance generalists | 0.124 | 0.124 | 0.125 | 0.127 |
| Unpaired t-test | p = 0.9305 | p = 0.323 | p = 0.255 | p = 0.197 |

***Table 2**: Average variable importance of the bioclimatic, lithology and topographic variables in the SDMs for glacial specialists versus generalist species and matching results of unpaired t-tests. Local bioclimate (Swiss scale) and global bioclimate (Europe scale) data are shown separately. The distinction between glacial specialists and generalists was drawn by varying the prevalence threshold of the glacial specialists in proglacial areas.*

A. Global bioclimate

| Prevalence | **T-index = 1** | **L_index = 5** | **W-index = 1** |
|---|---|---|---|
| Number of glacial specialist species | 87 | 136 | 191 |
| Number of generalist species | 282 | 233 | 178 |
| Average variable importance glacial specialists | 0.91 | 0.82 | 0.79 |
| Average variable importance generalists | 0.60 | 0.59 | 0.55 |
| Unpaired t-test | p<2.2e-16 | p = 2.444e-13 | 2.867e-13 |

B. Regional bioclimate

| Prevalence | **T-index = 1** | **L_index = 5** | **W-index = 1** |
|---|---|---|---|
| Number of glacial specialist species | 87 | 136 | 191 |
| Number of generalist species | 282 | 233 | 178 |
| Average variable importance glacial specialists | 0.048 | 0.060 | 0.064 |

| | | | |
|---|---|---|---|
| Average variable importance generalists | 0.074 | 0.073 | 0.072 |
| Unpaired t-test | p = 8.359e-05 | p = 0.054 | p = 0.269 |

C. Lithology

| Prevalence | T-index = 1 | L_index = 5 | W-index = 1 |
|---|---|---|---|
| Number of glacial specialist species | 87 | 136 | 191 |
| Number of generalist species | 282 | 233 | 178 |
| Average variable importance glacial specialists | 0.013 | 0.022 | 0.023 |
| Average variable importance generalists | 0.024 | 0.022 | 0.020 |
| Unpaired t-test | p = 0.0003 | p = 0.8352 | p = 0.35 |

D. Topography

| Prevalence | T-index = 1 | L_index = 5 | W-index = 1 |
|---|---|---|---|
| Number of glacial specialist species | 87 | 136 | 191 |
| Number of generalist species | 282 | 233 | 178 |
| Average variable importance glacial specialists | 0.118 | 0.119 | 0.117 |
| Average variable importance generalists | 0.125 | 0.126 | 0.130 |
| Unpaired t-test | p = 0.04248 | p = 0.018 | p = 5.043e-05 |

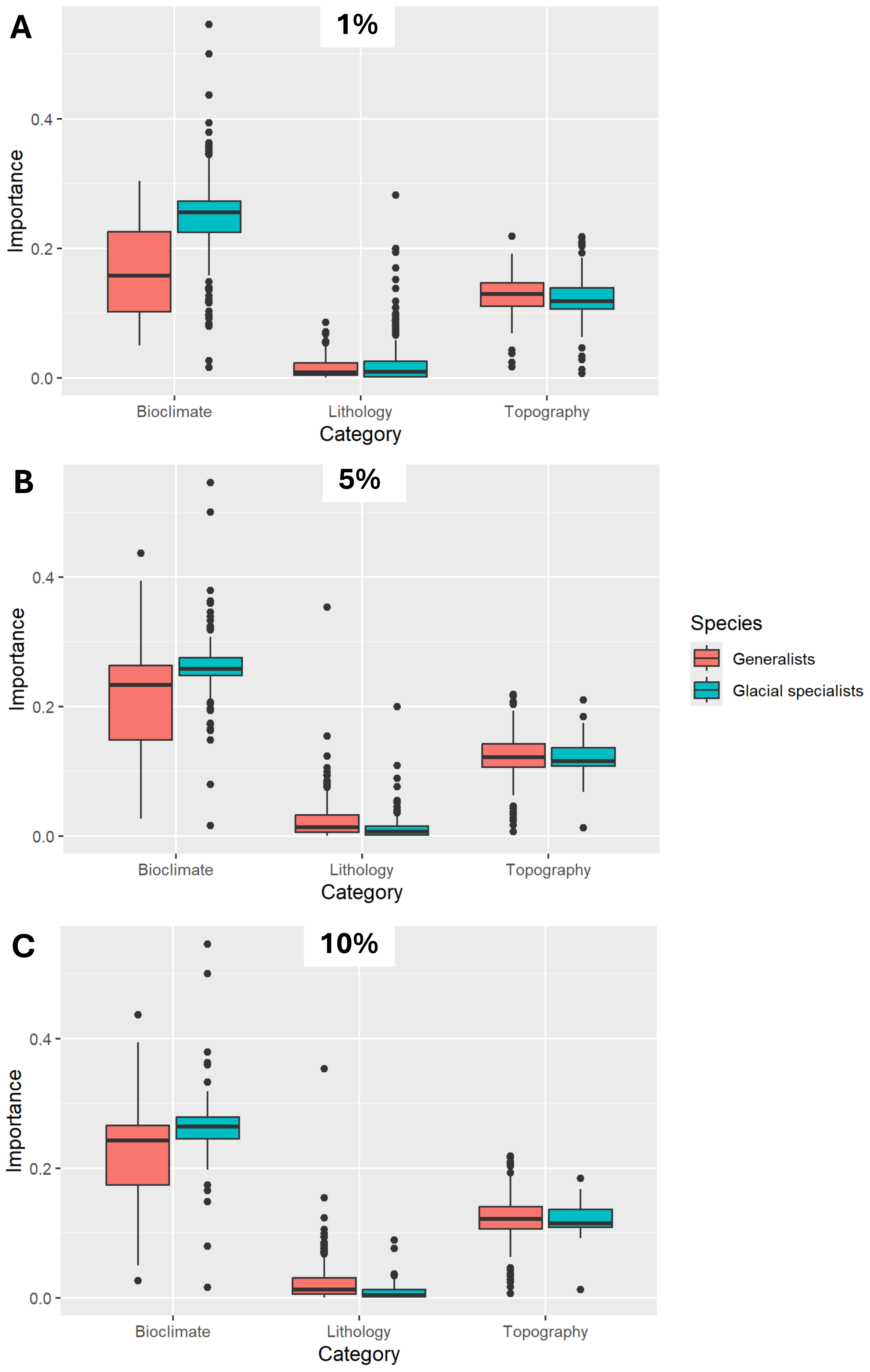
A
1%
Importance
0.0
0.2
0.4
Bioclimate
Lithology
Topography
Category
B
5%
Importance
0.0
0.2
0.4
Bioclimate
Lithology
Topography
Category
Species
Generalists
Glacial specialists
C
10%
Importance
0.0
0.2
0.4
Bioclimate
Lithology
Topography
Category

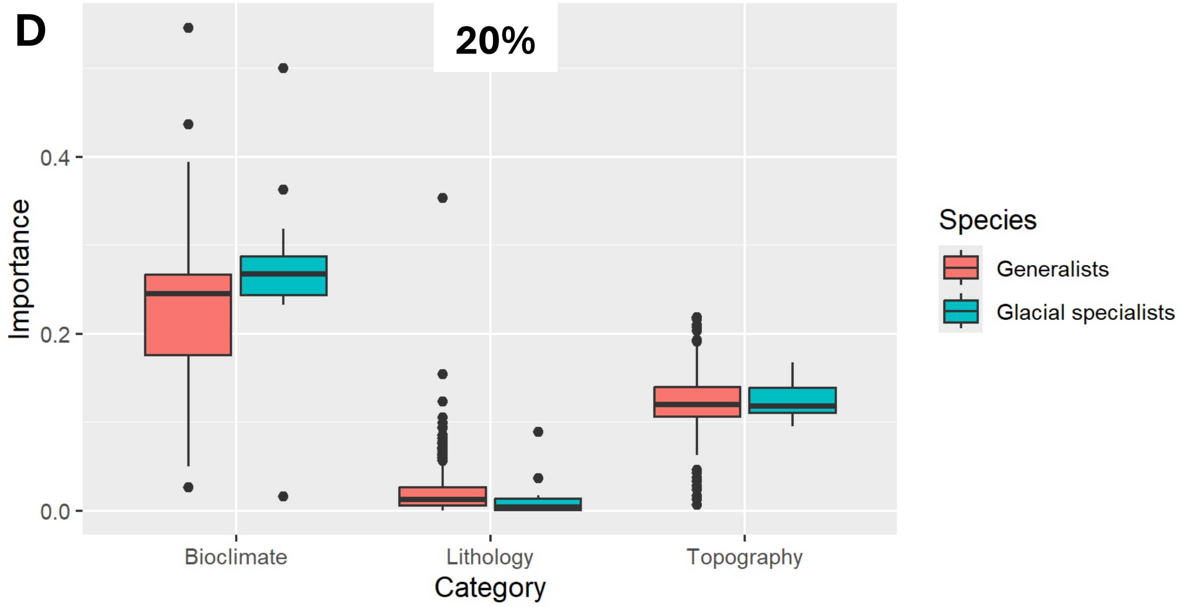


***Figure S1**: Average variable importance of the bioclimatic, lithology and topographic variables in the SDMs for glacial specialists versus generalist species. The distinction between glacial specialists and generalists was drawn by varying the prevalence threshold of the glacial specialists in proglacial areas (i.e., 1% prevalence = 1% or more of the total number of occurrence records were inside glacier forelands). The limits were: 1% (A), 5% (B), 10% (C) and 20% (D).*

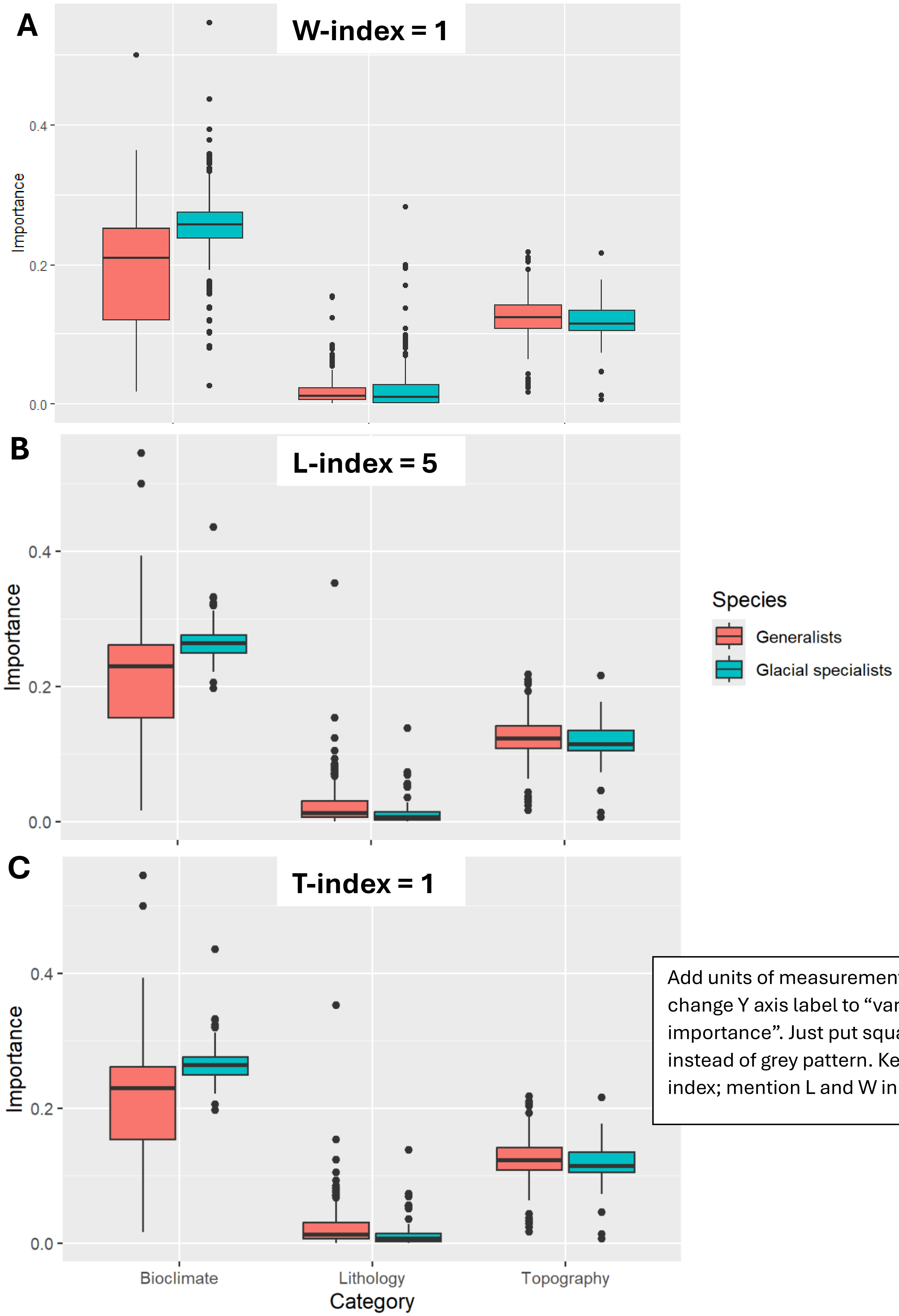


Add units of measurement ; change Y axis label to "variable importance". Just put square instead of grey pattern. Keep T-index; mention L and W in text.

***Figure S2****: Average variable importance of the bioclimatic, lithology and topographic variables in the SDMs for glacial specialists versus generalist species. The distinction between glacial specialists and generalists was drawn by alternating the Landolt index as follows: moisture variability (W) index equal to 1 (tolerates very little variation in moisture regimes) (A), light (L) index set to 5 (cannot tolerate shade) (B), T (temperature) index set to 1 (species belonging to alpine and/or nival habitats) (C).*

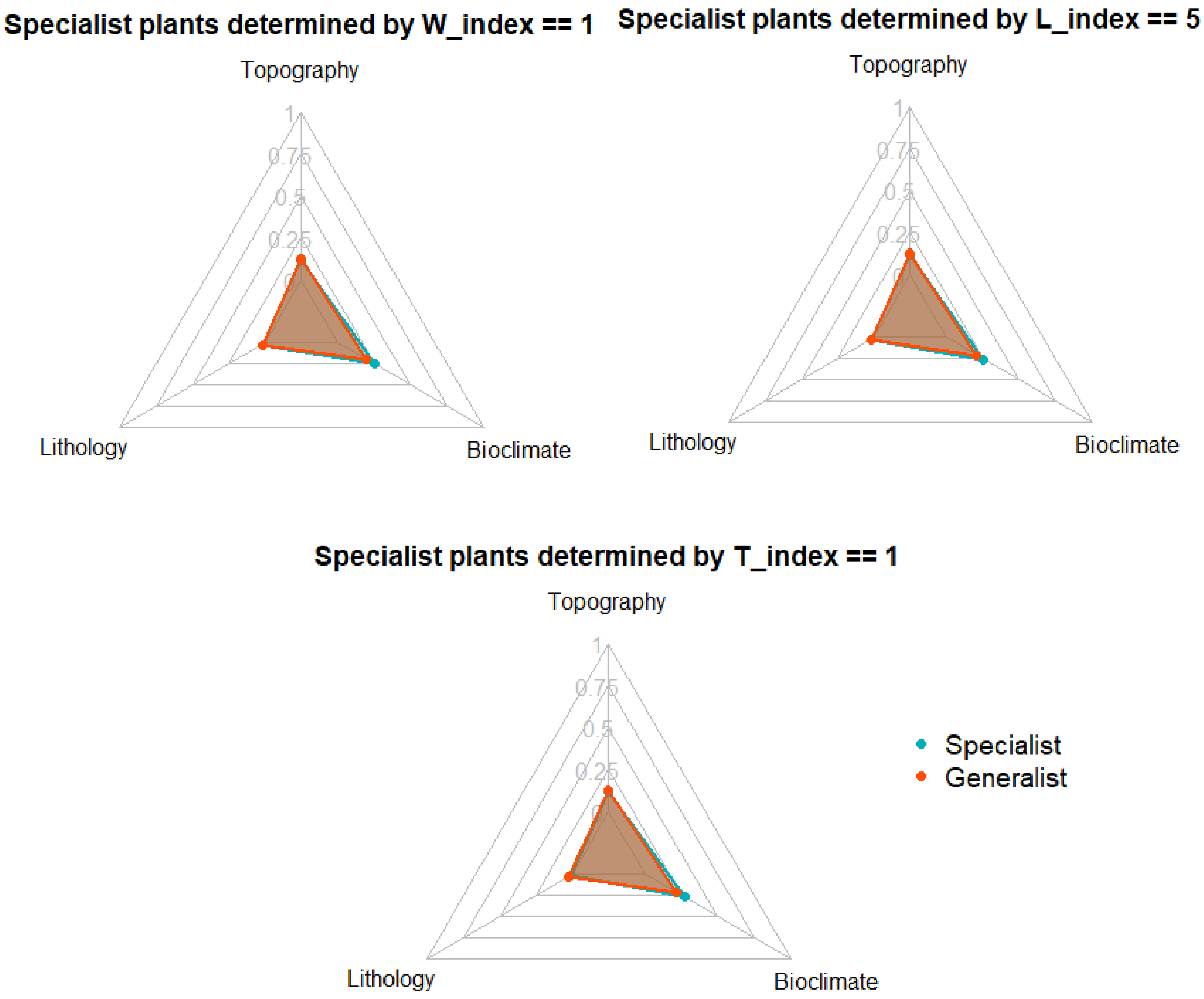


***Figure S3**: Average variable importance of the bioclimatic, lithology and topographic variables in the SDMs for glacial specialists versus generalist species. The distinction between glacial specialists and generalists was drawn by alternating the Landolt index as follows: moisture variability (W) index equal to 1 (tolerates very little variation in moisture regimes), light (L) index set to 5 (cannot tolerate shade), T (temperature) index set to 1 (species belonging to alpine and/or nival habitats).* ***Maybe move these to bin.***

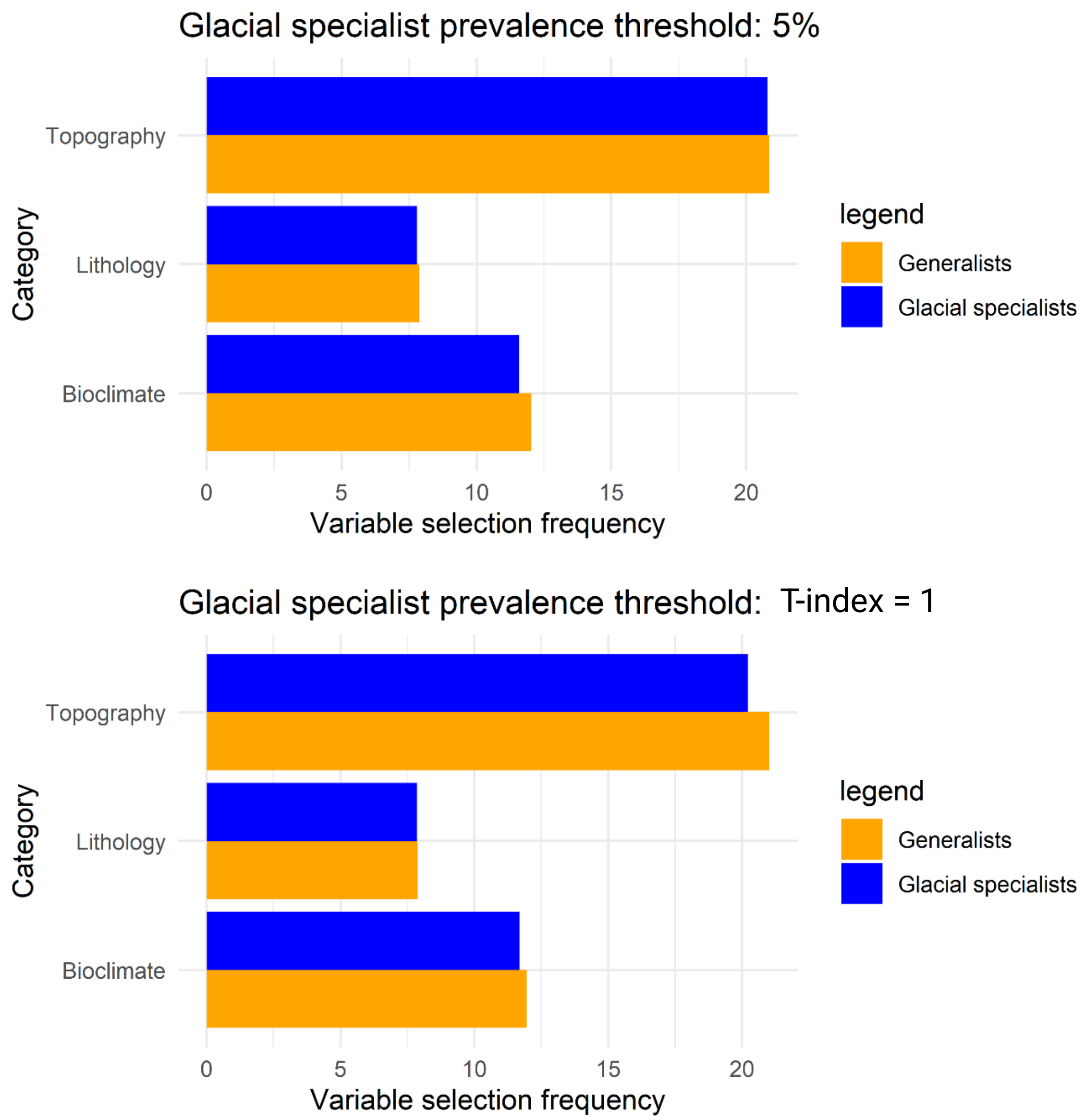


***Figure S5****: Average selection frequency of variables of the three categories (topography, bioclimate, lithology) by glacial specialist species versus generalist species. Distinctions between the species groups drawn by setting the prevalence threshold to 5% or setting the Landolt T-index value to 1 (meaning Swiss nival and/or glacial species).*
